# Global Profiling of Remodeled Subcellular Structures Due to Drug Treatment and Disease

**DOI:** 10.1101/2025.08.27.672480

**Authors:** Rachel A. Victor, Jesse J. Altemus, Michelle A. Lay, Sarah N. Shepherd, Valery F. Thompson, Mozani Zianti, Marco Padilla-Rodriguez, Austin Lipinski, Ghassan Mouneimne, Paul R. Langlais, Jacob C. Schwartz

## Abstract

Cellular biochemistry arises from various interactions between macromolecules, including proteins, nucleic acids, and lipids. These make up membrane-bound organelles, membrane-less compartments, and molecular assemblies and scaffolds. Changes due to stimuli or disease can significantly impact cell fate and metabolism. We recently reported our protocol combining crosslinking and size exclusion chromatography with mass spectrometry (SEC-MS). In this study, we explore global changes to subcellular structure in Ewing sarcoma cell lines or in response to drug treatment. Crosslinking-dependent differential enrichment across four human cell lines is seen for proteins involved in splicing, mitochondria function, and cell division. We confirmed changes to nucleoli structure. We also examine structures affected by a transcription inhibitor, flavopiridol. Following flavopiridol treatment, we observed changes to the levels of transcription and mRNA processing machinery present in large subcellular structures. Unexpected effects were also found, including structural changes to a cytoplasmic organelle, the peroxisome. Along with a reduction in peroxisome function, dissociation of peroxisome pore proteins PEX13 and PEX14 was detected by STORM microscopy. We conclude that SEC-MS combined with crosslinking is a valuable method to detect and quantify drug or disease effects on subcellular structures and may shed light on new aspects to mechanisms underlying their biologic outcomes.

**Graphical Abstract:** 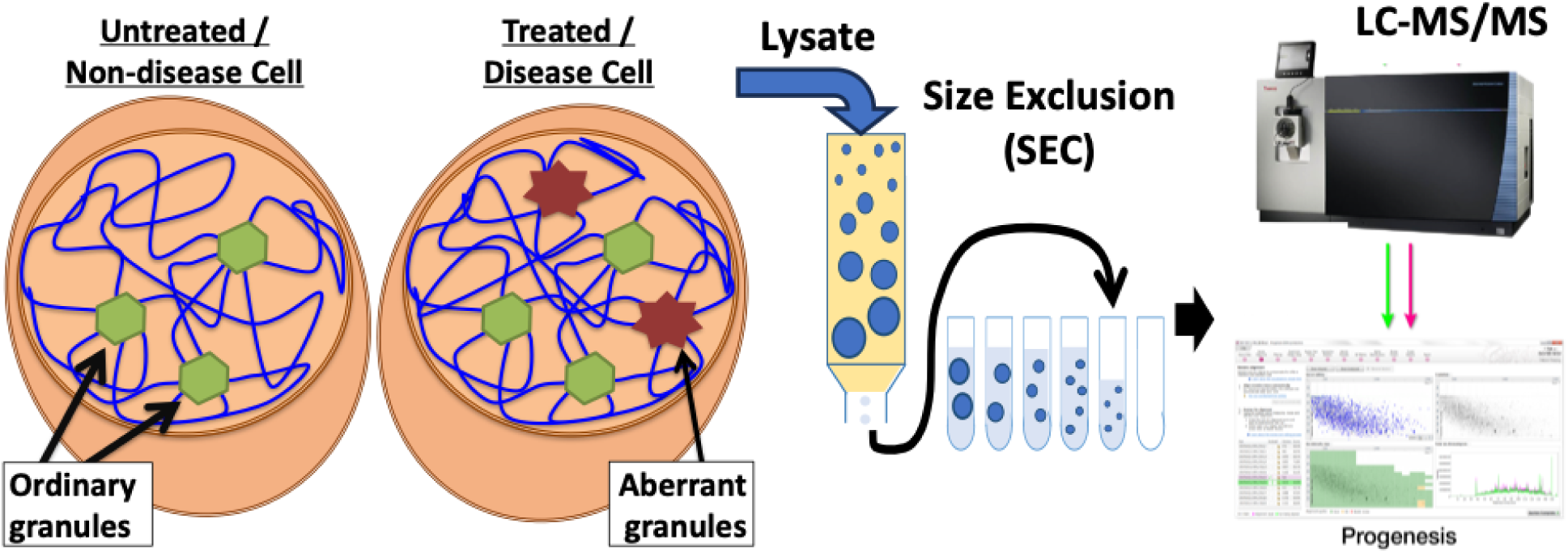

**Key Points:**

- Crosslinking plus SEC-MS shows global effects on subcellular structure in disease and treatment.
- Ewing sarcoma cell lines tested present distinct molecular composition in splicing, mitochondria, and nucleoli.
- A transcription inhibitor flavopiridol disrupts peroxisome function and protein import pores

## Introduction

In recent years, knowledge about basic cell biology and biochemistry has been reshaped by new understanding of phase-separated condensates^1, 2^. The primary mode of containment for cell biochemistry was once thought to be lipid membranes. Today, many complex signaling and metabolic mechanisms have been found to involve proteins containing low complexity domains with the ability to phase separate, a dynamic, membrane-less mechanism for compartmentalization^3, 4^. Phase-separated condensates, formed through multivalent interactions among proteins and nucleic acids, allow spatial and temporal control over processes such as gene expression, signal transduction, and stress response^4, 5^.

Compartmentalization makes it possible for biochemical processes, signaling pathways, or molecular mechanisms to occur within physiologic concentrations of proteins involved and in a controlled manner. For dynamic processes, like cell division or gene transcription, the physical properties of protein condensates offer rapid assembly, disassembly, and accommodation of macromolecular movement^1, 6^. Many proteins comprising subcellular structures and condensates are also be involved in human disease. Examples include the numerous proteinopathies found in amyloidosis, type II diabetes, and neurodegenerative diseases like Alzheimer’s disease, Parkinson’s disease, and amyotrophic lateral sclerosis (ALS)^7–9^. Many cancers are driven by fusion proteins that undergo phase separation, such as EWS-FLI1 in Ewing sarcoma, SS18-SSX1 or SS18-SSX2 in synovial sarcoma, PAX3-FOXO1 in rhabdomyosarcoma, and NUP98-HOXA9 in acute myeloid leukemia (AML)^10, 11^.

A challenge to the discovery of novel subcellular structures, especially condensates, is that they comprise of weak interactions that dissociate under cell lysis conditions. For this reason, structures such as Cajal bodies, p-bodies, and stress granules have been discovered by the punctate localization of functionally related proteins through cell microscopy methods^5, 12^. More recently, proteins with disordered domains able to phase separate in vitro are also employed as markers in the discovery of new subcellular bodies. This approach is labor intensive and in many recent examples, made possible through advanced microscopy methods not widely available to researchers^13^.

Finally, the local partitioning and structural organization within the cell alters drug mechanisms and accessibility to molecular targets^9, 14^. Structures that concentrate proteins and substrates can profoundly impact effective dose of drugs. Surprisingly, cell condensates have been found with the ability to sequester or exclude small molecule drugs^14^. Drugs are in development to treat disease by targeting condensates involved.

We sought a simplified unbiased method to quantitatively analyze the effects of molecular organization and structures on biology, disease, and drug development. Size Exclusion chromatography combined with mass spectrometry analysis (SEC-MS) is a tool to study the constitutional makeup of protein complexes and assemblies. SEC-MS combined with crosslinking has been well-established to provide finer resolution detail about binding interfaces and residue interactions^15^. SEC-MS can also be used as a discovery tool in cells to catalogue entire interactomes, and this approach is substantially improved when complexes are stabilized by chemical crosslinking^16, 17^. Examples of these approaches include correlation profiling with SILAC^18^, co-fractionation mass spectrometry^19^, *in vivo* crosslinking with co-fractionation^20^, and formaldehyde crosslinking with label-free SEC-MS^21, 22^.

While a powerful tool to discover interactomes, most fractionation SEC-MS protocols are limited to analyze protein complexes and assemblies only slightly larger than a ribosome, while the subcellular structures and membrane-less organelles described above are considerably larger. We have found previously developed SEC-MS protocols with formaldehyde crosslinking that can efficiently preserve protein interactions and used ELISA analysis to quantify proteins enriched in structures up to 400 nm or 10-times the diameter of a ribosome^23, 24^. We employed this method to study proteins individually that have significance in ALS and Ewing sarcoma^24, 25^. Recently, we combined our protocol with mass spectrometry to characterize recombinant expressed FUS protein and subcellular protein structures from *E. coli* lysates^26^. Here we test this SEC-MS protocol to identify differential crosslink-dependent associations across different cell lines and due to treatment by a cyclin-dependent kinase (CDK) inhibitor, flavopiridol.

## Methods

### Cell Culture

Cell lines A673 (ATCC, CRL-1598), SK-N-MC (ATCC, HTB-10), U2OS (ATCC, HTB-96), and HEK293T/17 (ATCC, CRL-11268) were obtained from ATCC and cultured at 5% CO_2_ and 37 °C. A673 and SK-N-MC were grown in DMEM supplemented with 10% fetal bovine serum (FBS). SK-N-MC media was also supplemented with sodium pyruvate (1 mM, ThermoFisher, cat. no. 11360070). Note: Throughout the document, HEK293T/17 is simplified to HEK293T. HEK293T were grown in DMEM supplemented with 5% FBS. U2OS cells were grown in McCoy’s 5A (Corning, cat. no. 10-050-CV) supplemented with 10% FBS. Flavopiridol used in cell treatments was purchased from Sigma-Aldrich (F3055-5MG).

### Crosslinking and Lysate Preparation

Cells were cultured in 15□cm dishes to confluence, washed with PBS, and incubated for 15 minutes in either PBS or 1% formaldehyde for crosslinking. Crosslinking was quenched by the addition of 1.5□mM glycine. Cells were harvested by scraping and washed once with PBS. Pellets were resuspended in five packed cell volumes of cold hypotonic lysis buffer (10□mM HEPES, pH 7.9, 1.5□mM MgCl□, 10□mM NaCl, 0.5□mM DTT, protease inhibitor cocktail, and 100 units Benzonase) and incubated on ice for 10 minutes. Cells were lysed by sonication using a Bioruptor Pico (Diagenode) for 30 cycles (30□sec on/30□sec off) at 4□°C. Lysates were cleared by centrifugation at >18,000□×□g for 30 minutes at 4□°C. Protein concentration was estimated by UV absorbance at 280□nm, and approximately 0.75□mg of total protein was loaded onto the size exclusion chromatography column.

### Size Exclusion Chromatography

A detailed protocol for size exclusion chromatography (SEC) methods has been previous published^23^. SEC was performed using Sepharose CL-2B resin (Cytiva, cat. no. 17014001) custom-packed into a 10 × 300 mm glass column, equilibrated with 2 column volumes of SEC running buffer (20 mM HEPES, pH 7.9, 100 mM NaCl, 0.2 mM EDTA, 5% glycerol, 6 M urea, 0.5 mM DTT, filter through 0.45 µm filter), and operated at a flow rate of 0.2 mL/min at 4 °C. Elution fractions were collected every 0.5 mL. Fractions were stored at -80 °C.

### In-Solution Tryptic Digest

SEC fractions were precipitated using 100% w/v trichloroacetic acid (TCA; Fisher Scientific, cat. no. bp555-500) at 4 parts to 1 part TCA and incubating 10 minutes on ice. The precipitates were centrifuged 16,000 x g for 5 minutes at 4 °C. After removing the supernatant, pellets were washed twice using 1 mL acetone followed by centrifugation at 16,000 xg for 5 minutes at 4 °C. Pellets were dried for 5-10 minutes at 95 °C, resuspended in 100 µL of 50 mM ammonium bicarbonate (pH 8.0; Thermofisher, cat. no. A643-500), and sonicated for 5 minutes. Reduction and alkylation of samples was performed by adding dithiothreitol (DTT) then incubated at 70 °C for 30 minutes, cooled to room temperature for 10 minutes, and incubated with 15 mM acrylamide for 30 minutes at room temperature while protected from light. The reaction was quenched with DTT with final concentration of 5 mM and incubated for 15 minutes in the dark. Proteins were first digested with 1□µg of Lys-C at 37□°C for 2–3 hours with shaking (300□rpm), followed by the addition of 50□µL of 50□mM ammonium bicarbonate and 2□µg of trypsin. Samples were incubated overnight at 37□°C with shaking (300□rpm). Trypsin digestion was quenched by the addition of 14.7□µL of 40% formic acid containing 1% heptafluorobutyric acid (FA/HFBA; final concentrations: 4% FA, 0.1% HFBA), followed by incubation for 10 minutes at room temperature. Peptides were desalted using Pierce™ Peptide Desalting Spin Columns (Thermo Scientific, cat. no. 89852) according to the manufacturer’s instructions, then dried by vacuum centrifugation. Dried peptides were resuspended in 20□µL of 0.1% formic acid (v/v), and peptide concentration was quantified using the Pierce™ Quantitative Colorimetric Peptide Assay Kit (Thermo Scientific, 23275) following the manufacturer’s protocol. A total of 350 ng of peptides was subjected to LC-MS/MS analysis.

### Mass Spectrometry

HPLC-ESI-MS/MS analysis was performed in positive ion mode on a Thermo Scientific Orbitrap Fusion Lumos Tribrid mass spectrometer equipped with an EASY-Spray source (Thermo Fisher Scientific)^27, 28^. Peptides were separated by nanoLC using a Thermo Scientific UltiMate 3000 RSLCnano system coupled to an EASY-Spray C18 column (50□cm × 75□µm i.d., PepMap RSLC C18, 2□µm particle size; Thermo Fisher Scientific, cat. no. ES803). Peptides were loaded at 0.300□µL/min for 15 minutes in Buffer A (0.1% formic acid in water), followed by a linear gradient from 1% to 34% Buffer B (0.1% formic acid in 80% acetonitrile) over 119 minutes at 0.220□µL/min. The gradient was ramped to 95% Buffer B over 4 minutes, held for 5 minutes at 0.250□µL/min, then returned to 1% Buffer B over 5 minutes and held for 10 minutes, for a total run time of 159 minutes. All solvents were LC-MS grade. Data acquisition was performed using Xcalibur software (version 2.3; Thermo Fisher Scientific).

### Label-Free Quantitative Proteomics

Label-free quantification based on ion intensity was performed using Progenesis QI for Proteomics software (version 2.4; Nonlinear Dynamics Ltd., Newcastle upon Tyne, UK) as previously described^29^. Raw MS data files (.raw) were imported and converted into two-dimensional ion maps (m/z vs. retention time), followed by automated alignment to a selected reference run. An aggregate dataset containing all detected features across all samples was generated, and only peptide ions with charge states of +2, +3, or +4 were selected for further analysis. Samples were grouped by condition (crosslinked vs. non-crosslinked), and peak lists of fragment ion spectra were exported in Mascot generic format (.mgf) for database searching. Searches were performed using Mascot (version 2.6; Matrix Science, London, UK) against Homo sapiens (SwissProt; 20,387 entries) protein databases. Search parameters included a precursor ion mass tolerance of 10 ppm, fragment ion mass tolerance of 0.5 Da, trypsin digestion with up to two missed cleavages, and variable modifications including methionine oxidation and phosphorylation of serine, threonine, and tyrosine. One ^13^C isotope was allowed. Mascot search results (.xml files) were imported back into Progenesis for peptide and protein assignment. Peptides with Mascot Ion Scores below 25 were excluded from quantification. Protein quantification was based on normalized precursor ion intensities, and only proteins identified by at least one unique peptide were considered for further analysis.

### Differential Protein Enrichment and Gene Set Enrichment Analysis

All proteins measured were exported from Progenesis. To calculate enrichment in a sample, the average crosslinked (XL) signal was divided by the average uncrosslinked (no XL) signal. Proteins were classified as enriched when log_2_ (XL/no XL) was > 0.75 and Anova(p) < 0.05. Differential enrichment for Ewing sarcoma or flavopiridol treated cells was also measured by the ratio of replicates for all Ewing sarcoma cell lines averaged and replicates for all non-Ewing sarcoma cell line averaged or flavopiridol replicates over vehicle-treated replicates. Functional annotation was performed using Database for Annotation, Visualization, and Integrated Discovery (DAVID) statistical overrepresentation tests (www.davidbioinformatics.nih.gov/summary.jsp) compared to reference lists for human genes and significance determined by Fisher’s exact test corrected for false discovery rate ^30^.

### Immunofluorescent Microscopy and Catalase Activity Analysis

Confocal images of nucleoli were acquired on a Nikon AX R Laser-Scanning confocal microscope using a Nikon 40x Plan Apo 0.95NA objective lens. Nucleoli were imaged with fibrillarin-specific antibody (Abcam, cat. no. ab4566), which was incubated overnight at 1:300 dilution. Confocal z-stacks was acquired using a 0.16µm/pixel resolution, with a 0.85 µm Zstep size covering the whole Z-range of the sample. Nikon NIS-Elements AR (v6.10) software with General Analysis 3 (GA3) was used for image processing and analysis. The following image processing functions were used to achieve accurate segmentation of the DAPI and Nucleoli channels: For DAPI, a low-pass filter (strength = 2px), rolling ball background subtraction (radius = 7 µm), and a gamma correction (gamma = 0.6) were applied. For Nucleoli, a low-pass filter (strength = 2px) and rolling ball background subtraction (radius = 3 µm) were applied. The Bright Spots Detection tool was used to threshold DAPI (diameter = 10 µm) and Nucleoli (diameter = 1 µm). To restrict the quantification of Nucleoli to the DAPI channel, the Binary AND function was used on the Nucleoli and DAPI binary layers to only measure the Nucleoli within the Nucleus. Cell nuclei that were touching the image border or undergoing mitosis were removed from the analysis. The number and area of each nucleoli were quantified per cell nucleus.

Peroxisomes were imaged in HEK293T cells were seeded at 1□×□10□cells per well in 8-well chamber slides (Corning, cat. no. 354108) and cultured for 24 hours to reach 70–80% confluence. Cells were transduced with BacMam 2.0 peroxisome-GFP (Thermo Fisher Scientific, cat. no. C10604) at a particle-per-cell (PPC) ratio of 30. After 24 hours, culture media (300□µL per well) was exchanged for fresh media with 1□µM flavopiridol or equal volume of vehicle (DMSO). One hour later, cells were aspirated and rinsed once with PBS then incubated with PBS containing 4% (v/v) paraformaldehyde (PFA) for 20 minutes at room temperature. Cells were gently rinsed with PBS and incubated in PBS with 100□µL of Phalloidin-iFluor™ 555 (Cayman Chemical, cat. no. 20552) and 1% (w/v) BSA for 30 minutes at room temperature. Coverslips were mounted using 25□µL of Vectashield® Antifade Mounting Medium with DAPI (Vector Laboratories, cat. no. H-1200-10). Confocal images were acquired on an Olympus Fluoview FV1200 microscope. Peroxisomes were quantified using the Particle Analyzer plugin in Fiji/ImageJ, with size thresholds set to 0.1–1□µm to exclude background signal^31, 32^.

Catalase activity was measured for HEK293T cells seeded in 6-well dishes at 1 x 10^6^ per well, allowed to grow 24 hours, then treated for 1 hour by 1 µM flavopiridol or DMSO vehicle. Catalase activity assays were performed using the CAT Activity Assay Kit (Elabscience, cat. no. E-BC-K031) according to the manufacturer’s instructions.

### STORM Microscopy

For STORM imaging, No. 1.5 glass-bottom poly-D-lysine–coated 35□mm dishes (MatTek Corporation, cat. no. P35GC-1.5-14-C.S) were rinsed with 100% ethanol, air-dried, and seeded with 2□×□10□HEK293T cells per dish in complete DMEM supplemented with 5% FBS. After 24 hours, when cells reached approximately 50–70% confluence, they were treated with either 1□µM flavopiridol (diluted from a 1□mM DMSO stock) or DMSO vehicle control for 1 hour at 37□°C in a humidified incubator with 5% CO□. Cells were then washed twice with PBS and fixed in 4% paraformaldehyde (PFA) in PBS for 20 minutes at room temperature (RT), followed by three 5-minute washes in PBS. Permeabilization was performed in 0.1% Triton X-100 for 30 minutes at RT, followed by direct incubation (no wash) in blocking buffer (1% FBS and 1% BSA in PBS) for 30 minutes at RT.

Primary antibodies diluted in the same blocking buffer were applied without intermediate washing and incubated for 1 hour at RT: anti-PEX13 (mouse monoclonal, clone D-5; Santa Cruz Biotechnology, cat. no. sc-271477) at 1:1000, and anti-PEX14 (rabbit polyclonal; Thermo Fisher Scientific, cat. no. PA5-78103) at 1:5000. Cells were then washed three times with washing buffer (0.2% BSA and 0.05% Triton X-100 in PBS), followed by 1-hour incubation at room temperature with secondary antibodies diluted 1:1000 in blocking buffer: either goat anti-mouse IgG Alexa Fluor 647 (Thermo Fisher Scientific, cat. no. A-21236) or donkey anti-rabbit IgG Alexa Fluor 488 (Thermo Fisher Scientific, cat. no. A-21206). After staining, cells were washed three times with washing buffer, then three additional times with PBS.

A second fixation step was performed using 4% PFA in PBS for 10 minutes at room temperature, followed by three PBS washes. Dishes were stored in PBS supplemented with 0.02% sodium azide, sealed with parafilm, and protected from light at 4□°C until imaging. Immediately before STORM imaging, dishes were rinsed three times with PBS and once with filtered Milli-Q water. Imaging buffer was prepared by mixing 7□µL GLOX solution, 7□µL 2-mercaptoethanol, and 690□µL Buffer B. Cells were mounted in Attofluor cell chambers, and STORM images were acquired following image protocols^33^.

Imaging was performed using 3D N-STORM 5.0 system with astigmatic lens (Nikon), with a CFI HP Apochromat 100XAC TIRF 1.49 NA objective (Nikon), LU-NV 405 nm (20 mW), 488 nm (70 mW), 561 nm (70 mW), and 640 nm (125 mW) laser with power measured at the fiber tip (Nikon), iXon Ultra 897 (Andor), and a motorized stage. For image acquisition, cells were located within the TIRF module, and a 40,000 frame imaging cycle was used for STORM imaging in 488-, 561-, and 640-nm wavelengths. Distances were measured between PEX13 (647 nm) and the nearest PEX14 (488 nm) across in 22 DMSO-treated and 9 flavopiridol-treated cells.

## Results

### Enrichment by crosslinking identifies subcellular structures in human cells

We sought to determine enrichment of human subcellular structures through crosslinking using SEC-MS. We chose cell lines representing two genetic backgrounds: Ewing sarcoma (A673 and SK-N-MC) and non-Ewing sarcoma (HEK293T and U2OS). Following our previously published protocol, cells were grown to confluency, harvested, and divided into samples crosslinked with formaldehyde or uncrosslinked (**Figure 1A**). After sonication and nuclease treatment, lysates were separated by SEC and fractions pooled into “Large” (150 – 450 nm), “Medium” (50 – 150 nm), and “Small” (< 50 nm).

**Figure 1.**
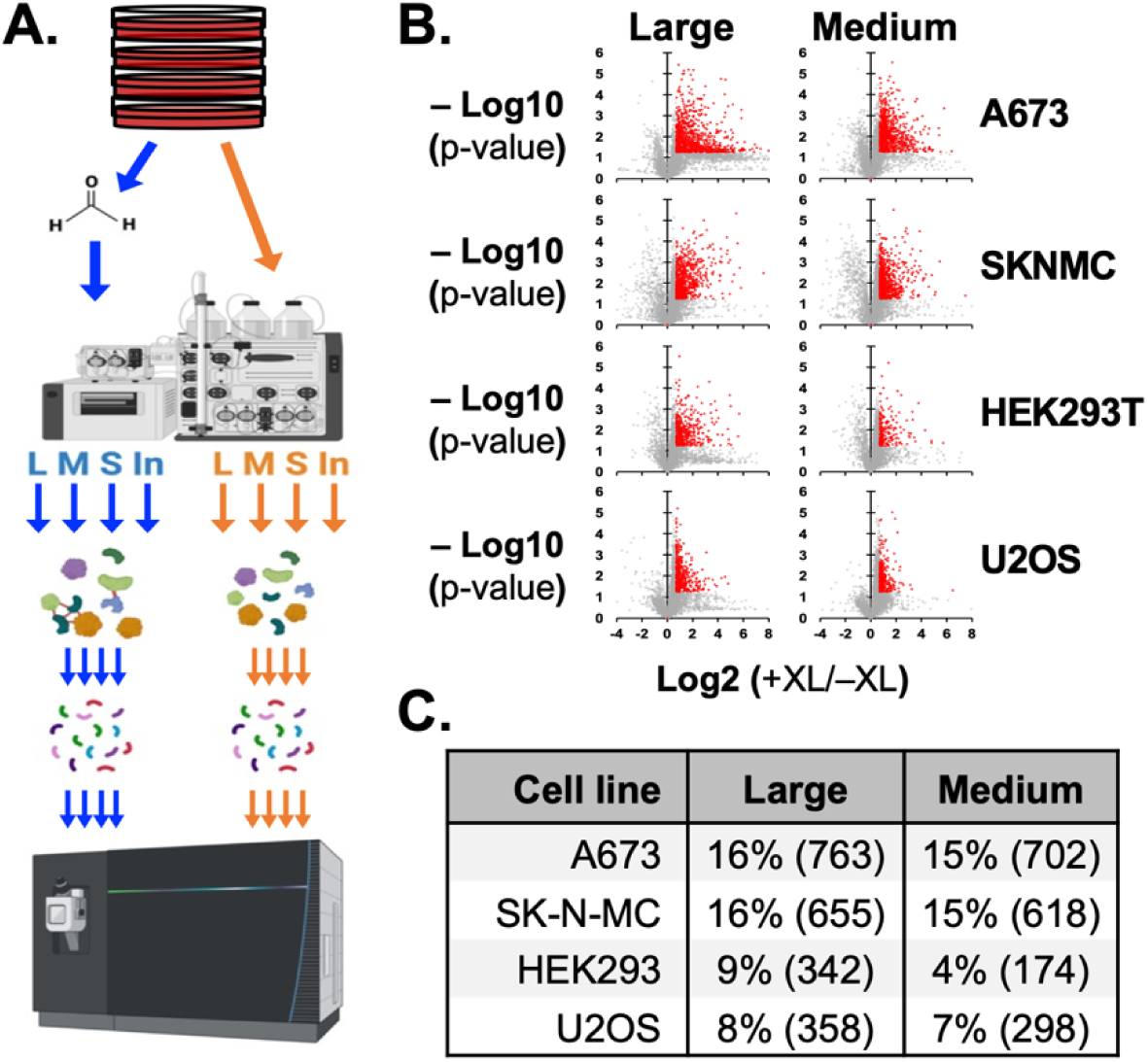
SEC-MS of human cell lysates. (**A**) Workflow to analyze protein particles through crosslinking, size-exclusion chromatography, and LS-MS/MS. (**B**) Volcano plots show enrichment of proteins quantified in large and medium particle pools across 4 cell lines. Red indicates proteins enriched more than 1.7-fold with p < 0.05, student T-test assuming equal variances. (**C**) Table shows the number of proteins enriched in cell line and pool and the % of proteins enriched among all proteins quantified in that pool.

Protein pools were precipitated and resuspended to perform protease digestion and then LC/MS-MS. The mass spectrometry-based proteomics analysis yielded high-quality data, as assessed by several key metrics. The average mass error across all spectra was below 5 ppm, indicating high mass accuracy. The average coefficient of variation for quantified proteins was 18.6 ± 6.1% for replicates with and without crosslinking for the 4 pools and each cell lines (**Supplemental Table 1**).

Fold enrichment by crosslinking and significance was determined for each cell line in triplicate and each pool and input sample. For the A673 and SK-N-MC Ewing sarcoma cell lines, 763 and 655 proteins were significantly enriched by >1.7-fold in large particles, respectively, and 702 and 618 in medium particles (**Figure 1B-C**). For non-Ewing HEK293T and U2OS cell lines, 342 and 358 proteins were enriched in large particles and 174 and 298 proteins respectively in medium particles. We also noted that a larger fraction of all proteins detected in the large pool of Ewing sarcoma cell lines were enriched by crosslinking, 16% for A673 and SK-N-MC, relative to non-Ewing cell lines, 9% for HEK293T and 8% for U2OS.

We first investigated proteins enriched by crosslinking in all cell lines. We determined that 340 proteins in the large particle pools and 175 proteins in medium particles were significantly enriched across both genetic backgrounds (**Supplement Figure 1A-B**). We next performed a broad functional annotation clustering of the combined 494 proteins significantly enriched in both large and medium particles sizes and genetic backgrounds using the DAVID bioinformatics tool (**Figure 2A**). The top 10 clusters included GO terms for translation and ribosomal proteins. Mitochondrial protein associations were also among the most significantly enriched. Consistent with general properties of biomolecular condensates, RNA-binding protein and unfolded protein binding or proteosome associations were among the enriched clusters (**Figure 2A**, **Supplemental Table 2**).

**Figure 2.**
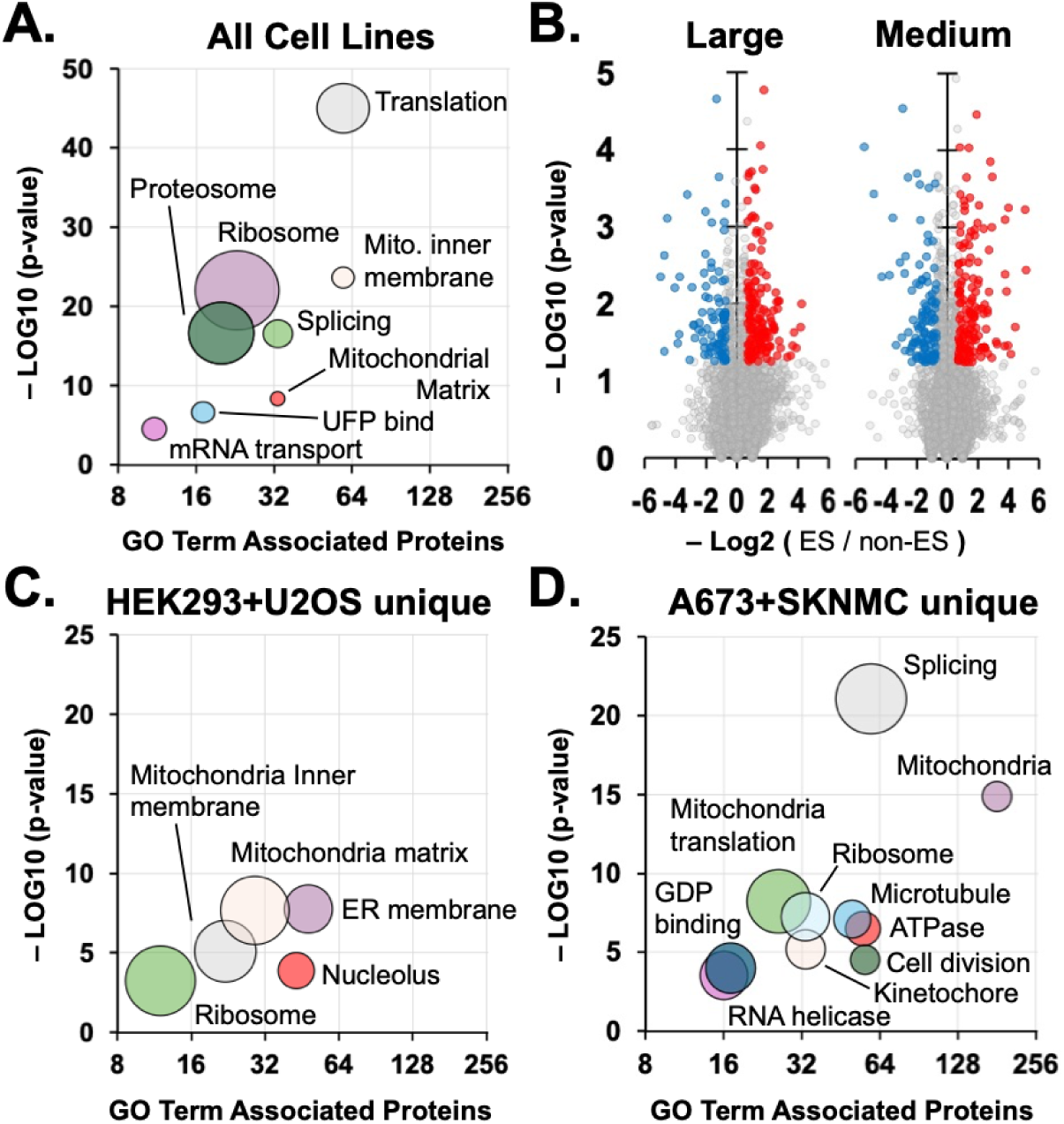
Functional Annotation Cluster analysis of proteins enriched in Ewing sarcoma derived cell lines, A673+SK-N-MC, and non-Ewing cell lines, HEK293T+U2OS. (**A**) Bubble plot show selected functional annotation clusters and GO terms for proteins enriched by crosslinking across all cell lines. Diameters indicate fold enrichment of associated proteins. (**B**) Volcano plots compare relative protein enrichment that is significantly higher in Ewing sarcoma (red) or non-Ewing sarcoma (blue) cells. (**C**) Plot shows the most significant clusters associated with proteins uniquely enriched among large and medium particles of non-Ewing cell lines. (**D**) Plot shows th same for Ewing sarcoma derived cell lines.

We hypothesized that cellular component associations should identify known subcellular structures expected to be present regardless of cell type or background and we inspected enriched proteins and their functional associated structures (e.g. GO Cellular Component) or interactomes (e.g. INTACT) using the DAVID bioinformatics tool. We noted enrichment for the membrane bound structures that include focal adhesions, nuclear pores, and nuclear matrix (**Table 1**). Mitochondrion proteins enriched included membrane bound structures and nucleoid proteins.

**Table 1.** Cellular component term associations among 494 proteins enriched in large and medium particles across 4 human cell lines.

| GO Cellular Component Associations (FC>2.5) | Count | Fold Enrich | -log(p) | Related GO Cellular Component Associations (-log(p-value)) |
| --- | --- | --- | --- | --- |
| GO:0022626~cytosolic ribosome | 43 | 19.7 | 43.0 | GO:0022625-cytosolic large ribosomal subunit (25.5); GO:0022627-cytosolic small ribosomal subunit (22.0); GO:0005840-ribosome (18.8); GO:0015935-small ribosomal subunit (9.4) |
| GO:1990904~ribonucleoprotein complex | 43 | 8.8 | 26.5 | GO:0010494-cytoplasmic stress granule (5.6) |
| GO:0005739~mitochondrion | 97 | 2.8 | 19.9 | GO:0005743-mitochondrial inner membrane (23.6); GO:0005759-mitochondrial matrix (8.3); GO:0042645-mitochondrial nucleoid (7.9); GO:0005753-mitochondrial proton-transporting ATP synthase complex (5.0) |
| GO:0000502~proteasome complex | 20 | 14.9 | 16.6 | GO:0005839-proteasome core complex (16.5); GO:0019773-proteasome core complex, alpha-subunit complex (7.2); GO:0019774-proteasome core complex, beta-subunit complex (6.9) |
| GO:0005925~focal adhesion | 46 | 4.6 | 16.5 | GO:0045296-cadherin binding (8.8) |
| GO:0005730~nucleolus | 82 | 2.5 | 13.8 | GO:0032040-small-subunit processome (16.5); GO:0007000-nucleolus organization (2.2) |
| GO:0005681~spliceosomal complex | 20 | 6.5 | 9.7 | GO:0071013-catalytic step 2 spliceosome (9.8); GO:0000398-mRNA splicing, via spliceosome (16.5); GO:0008380-RNA splicing (13.9); GO:0005686-U2 snRNP (4.3) |
| GO:0005694~chromosome | 23 | 3.9 | 6.9 | GO:0003676-chromatin binding (3.1); GO:000776-kinetochore (2.7); GO:0007064-mitotic sister chromatid cohesion (1.3) |
| GO:0005643~nuclear pore | 13 | 5.8 | 5.6 | GO:0006913-nucleocytoplasmic transport (3.8); GO:0031965-nuclear membrane (2.1) |
| GO:0016607~nuclear speck | 28 | 2.8 | 5.5 |  |
| GO:0016363~nuclear matrix | 14 | 4.6 | 5.0 |  |

Nuclear structures enriched included the nucleolus, ribonucleoprotein complexes, spliceosomal complexes, and nuclear speckles, which are all notable as condensates with liquid-like properties. The remaining cell component associations were cytosolic ribosomes, proteasome complex, and chromosomal proteins (**Table 1**). For protein associations by interactome (INTACT), the same cell component associations were identified and two more subcellular organelles, the Cajal body and P-body (**Supplemental Table 2**).

We investigated whether large or medium pools from 2 or more cell lines were uniquely enriched for protein associations. For example, U1 snRNP proteins were enriched only in large pools (N = 6, p = 10^-3^), not Medium (**Supplemental Table 2**). However, U2 associated snRNP and spliceosome complexes were enriched in both large and medium pools. Similarly, associations for the proteosome (N = 21, p < 10^-18^), protein folding (N = 20, p < 10^-7^), and chaperones (N = 9, p < 10^-4^) were unique to the large pool. Enrichment of transcription machinery that includes subunits of RNA Pol II, mediator complex, and initiation factors, was found in medium-sized particles (N = 8, p = 0.02), while transcription machinery enriched in the large pool were those associated with elongation (N = 5, p = 0.02). Nuclear speckle proteins were also uniquely enriched among medium-sized particle proteins (N = 16, p = 10^-4^).

### SEC-MS analysis of two Ewing sarcoma cell lines indicates defects in nucleoli formation

Finding that proteins enriched in large or medium particles across 4 human cell lines showed consistent and predicted associations with known structures, we asked if SEC-MS with crosslinking identified differential enrichment of proteins in large and medium particles that could indicate differences in structural makeup of the two Ewing sarcoma cell lines and two non-Ewing sarcoma cell lines.

From large and medium particle pools collected through SEC, we identified enrichment for 1437 proteins that was unique among two Ewing sarcoma cell lines and 278 unique to non-Ewing sarcoma lines, termed “A673+SK-N-MC unique” and “HEK293T+U2OS unique,” respectively (**Figure 2B**). The 494 proteins enriched in both groups were labeled as “shared.” We performed functional annotation cluster analysis on HEK293T+U2OS unique and found 5 clusters with >10 proteins associated in GO terms (**Figure 2C**). The HEK293T+U2OS unique proteins were most distinguished for enriched in large and medium particles is the representation of ER membrane and nucleolar proteins. On the other hand, A673+SK-N-MC unique proteins were most enriched in large and medium particles for spliceosomal and mitochondria proteins, followed by translation and cell division related proteins, microtubule, and RNA helicases (**Figure 2D**).

Further comparison by clusters highlighted associations in common and striking differences between proteins enriched shared by both, in A673+SK-N-MC only, or in HEK293T+U2OS only across large and medium particles. For example, clusters revealed across all groups, 59 shared proteins formed a significant association with translation (p < 10^-45^, GO:0006412), as well as 30 A673+SK-N-MC unique (p = 10^-4^) and 8 HEK293T+U2OS unique (p = 0.02) proteins (**Table 2**). Similarly, 97 shared mitochondrion proteins (p < 10^-19^, GO:0005739) were enriched, as were 181 A673+SK-N-MC unique (p < 10^-14^) and 53 HEK293T+U2OS unique (p < 10^-10^) proteins (**Table 2**).

**Table 2.**
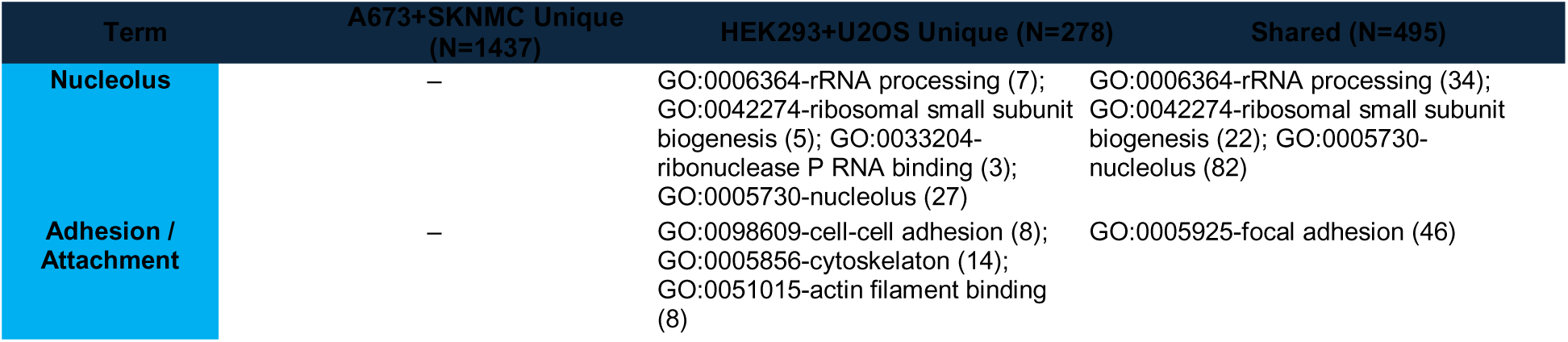
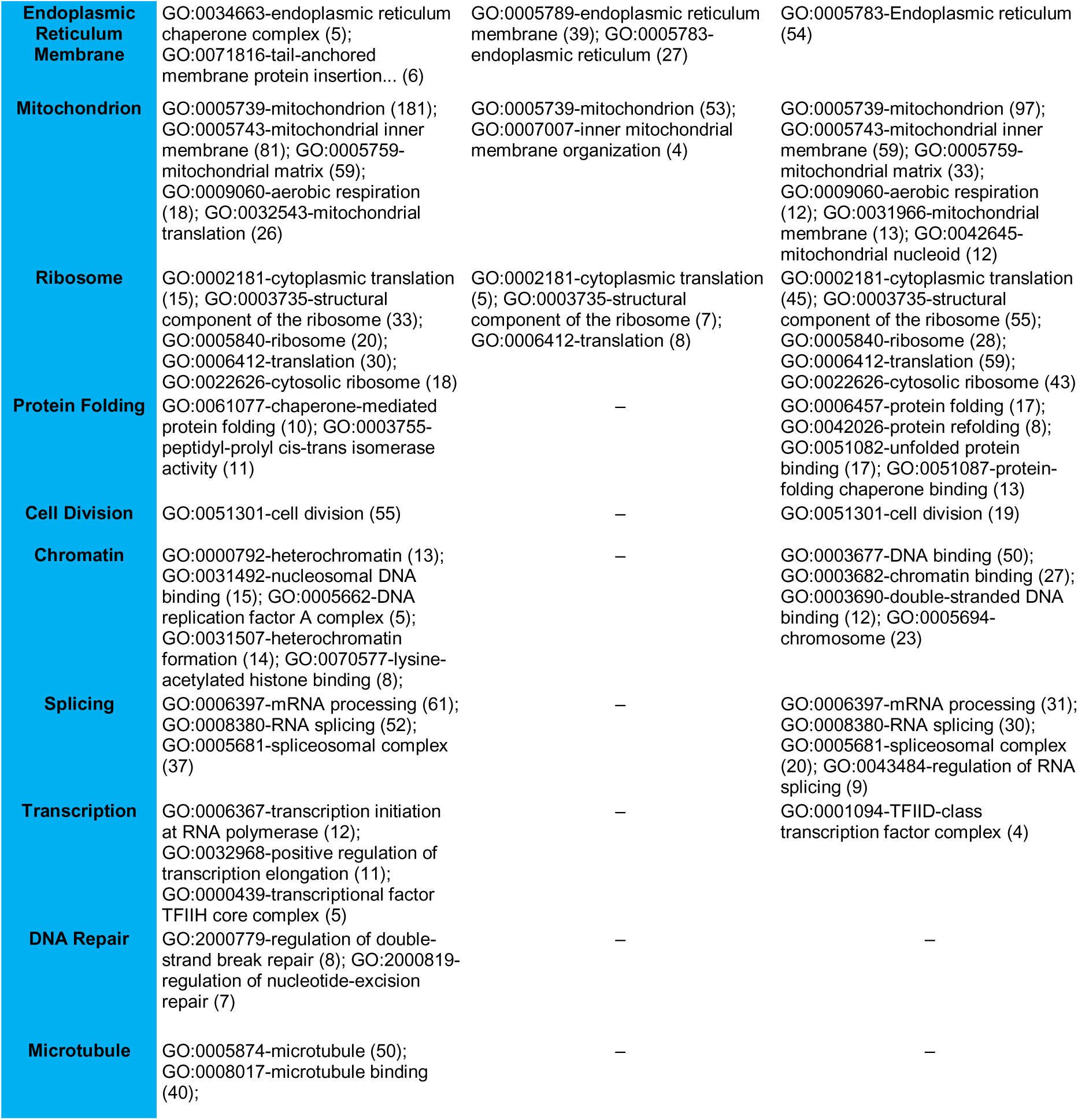
Comparison of functional annotation clustered GO terms identified for proteins uniquely enriched in Ewing sarcoma cell lines (A673+SK-N-MC), non-Ewing sarcoma cell lines (HEK293T+U2OS), or both (Shared).

A functional association cluster for protein folding was found for shared and A673+SK-N-MC unique but not HEK293T+U2OS unique proteins. Shared proteins were enriched for terms including protein folding (N = 17, p < 10^-5^) and protein-folding chaperone binding (N = 13, p = 10^-5^) and A673+SK-N-MC unique proteins for chaperone-mediated protein folding factors (N = 10, p < 10^-2^) and peptidyl-prolyl cis-trans isomerase activity (N = 11, p < 10^-2^) (**Table 2**). The associations with cell division and mRNA processing were made by twice or more the number of A673+SK-N-MC unique proteins compared to shared proteins. Chromatin and transcription associations were notably greater for A673+SK-N-MC unique relative to shared proteins. Lastly, significant associations with DNA repair and microtubules were found only for A673+SK-N-MC unique proteins (**Table 2**).

Finally, we chose nucleolar proteins as a functional association among shared and HEK293T+U2OS proteins but not detected in proteins enriched uniquely in Ewing sarcoma cell lines. For this purpose, nucleoli are imaged using antibody specific to fibrillarin for A673 cells, HEK293T cells, and HEK293T cells transfected by plasmid to express EWS-FLI1 (**Figure 3A**). The area of nucleoli for A673, HEK293T, and HEK293T(+EWS-FLI1) did not differ significantly (**Figure 3B**). The number of nucleoli in A673 cells was significantly higher than for HEK293T cells (N = 4, p < 0.01) (**Figure 3C**). HEK293T cells expressing EWS-FLI1 also contained more nucleoli per nucleus (N = 6, p = 0.03) and was not significantly different from A673 cells (N = 6, p = 0.1).

**Figure 3.**
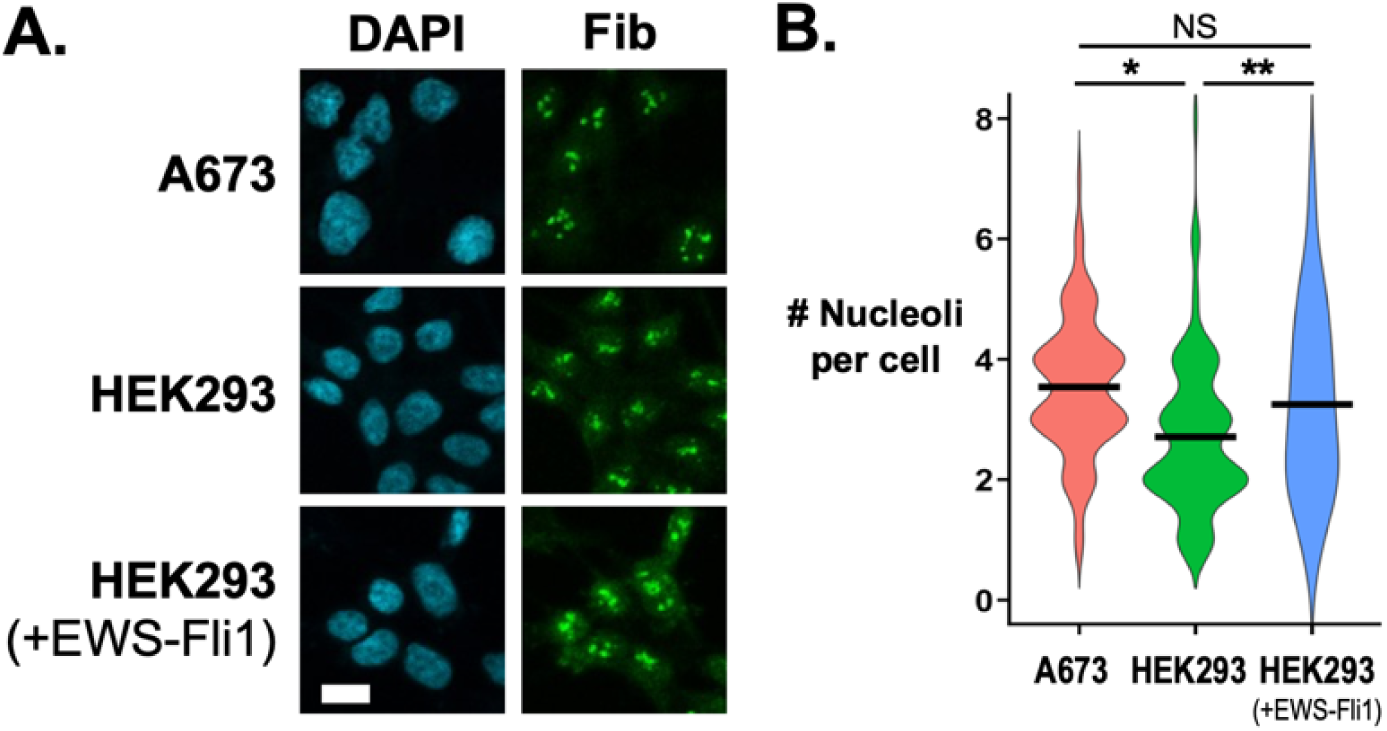
Comparison of nucleolar structure in A673 and HEK293T cells. (**A**) DAPI stain of DNA and fibrillin stain of nucleoli are shown for A673 cells, HEK293T cells, and HEK293T cells overexpressing EWS-FLI1 from plasmid. Scale bar = 10 µm. Images shown are representative of between 4 and 6 biologic replicates. (**B**) Nucleoli per cell was counted for A673 (n = 236), HEK293T (n = 473), and HEK293T expressing EWS-FLI1 (n = 516). Bars indicate median for the set. Significance was measured by Student T-Test: * p < 0.05; ** p < 0.01; NS = not significant.

### SEC-MS analysis of blocking transcription reveals defect in peroxisome protein transport

We next explored further whether differential enrichment by crosslinking indicated a subcellular structure change that is visible to orthogonal approaches. We investigated the response of subcellular structures in a single cell line to inhibition of RNA Pol II transcription with 1 μM flavopiridol for 1 hour. This protocol has been previously used to produce a near complete block of mRNA production, and the short incubation period minimizes protein reduction that could impact subcellular structure^24^.

In flavopiridol treated HEK293T cells, we detected 574 proteins significantly enriched in large particles and 504 in medium particles. From the DMSO treated cells, 643 large and 961 medium particle proteins were identified. The number of HEK293T proteins enriched in large or medium particles was approximately double that in our previous experiment (**Figure 1C**). We reason this to be a result of an increased protein yield during lysis and SEC.

We measured the change in enrichment of proteins in large and medium particles for flavopiridol relative to DMSO treated samples (**Figure 4A**). We found 73 proteins with significantly increased enrichment (≥ 1.5-fold, p < 0.05 student t-test) in large particles and 102 proteins for medium particles, making a total of 169 unique proteins combined. Protein enrichment was decreased for 93 proteins in large and 146 in medium particles, or 236 combined (**Figure 4A**).

**Figure 4.**
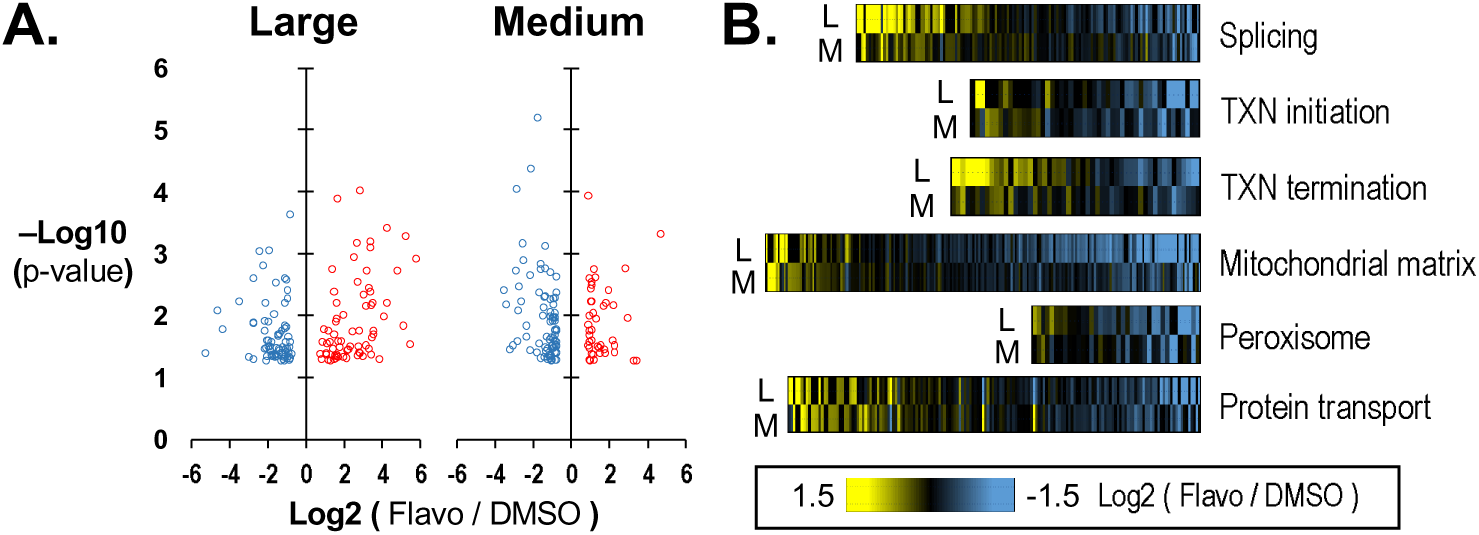
Differential enrichment of proteins in cells treated with flavopiridol or DMSO. (**A**) Volcano plots show differential enrichment by crosslinking for large and medium particle proteins measured in cells treated with flavopiridol relative to vehicle only, DMSO. (**B**) Heat plots show the enrichment of proteins associated with selected GO terms representing transcription, RNA processing, mitochondrial function, peroxisomes, and protein transport.

The most significant associations for proteins increased in enrichment for large or medium particles included protein transport (p < 10^-3^), mRNA splicing (p < 10^-2^), nuclear speckles (p < 10^-4^), and unfolded protein binding (p < 10^-3^) (**Figure 4B**, **Supplemental Figure 2A**). Terms from proteins decreased in enrichment included mitochondrial matrix (p < 10^-9^), endoplasmic reticulum (p < 10^-3^), peroxisomal membrane (p < 10^-3^), and DNA repair (p < 0.02) (**Figure 4B**, **Supplemental Figure 2B**). Because the target of flavopiridol is the P-TEFb kinase, we manually inspected proteins associated with transcription. We noted that most changes in enrichment among proteins involved in transcription initiation were decreased with flavopiridol treatment (**Figure 4B**). This included a 9-fold decrease in enrichment for the cyclin T1 subunit of P-TEFb among medium particle proteins (p < 0.01). The effect on transcription initiation factors was consistent with the response of RNA Pol II granules to flavopiridol in our related previous study^24^. In contrast, most changes in protein enrichment associated with transcription termination were increased in the large particle pools (**Figure 4B**, **Supplemental Table 2**).

### Peroxisome pore proteins are disassembled by flavopiridol treatment

While functional annotation of proteins found increased protein transport proteins in large or medium particles after flavopiridol treatment, this was not true of those associated with the peroxisome organelle (**Figure 4B**). Moreover, while flavopiridol effects on mRNA production was expected, there was not obvious reason to predict such immediate changes to peroxisome protein assemblies. Peroxisomes are small, membrane-bound organelles that play a key role in breaking down fatty acids and detoxifying reactive oxidative species^34, 35^. While peroxisomes produce hydrogen peroxide by metabolizing fatty acids, they contain catalase enzyme to convert hydrogen peroxide and prevent cell damage. We reasoned that validation of flavopiridol effects on peroxisome function would test the utility of this approach as a tool for discovery.

We first measured number of peroxisomes per cell by expressing green fluorescent protein (GFP) fused to a peroxisome targeting sequence (PTS1). HEK293T cells were transduced by PTS1 expressing BacMam 2.0 for 24 hours prior to treatment with flavopiridol or DMSO for 1 hour, then analyzed by confocal microscopy (**Figure 5A**). Treatment by flavopiridol did not change the number of peroxisomes observed through PTS1, except shift the apparent ratio of large and small diameter peroxisomes (**Figure 5B, Supplemental Figure 3**). However, we tested function of peroxisomes and found a significant reduction in catalase activity, indicating that peroxisome function was diminished by flavopiridol (**Figure 5C**).

**Figure 5.**
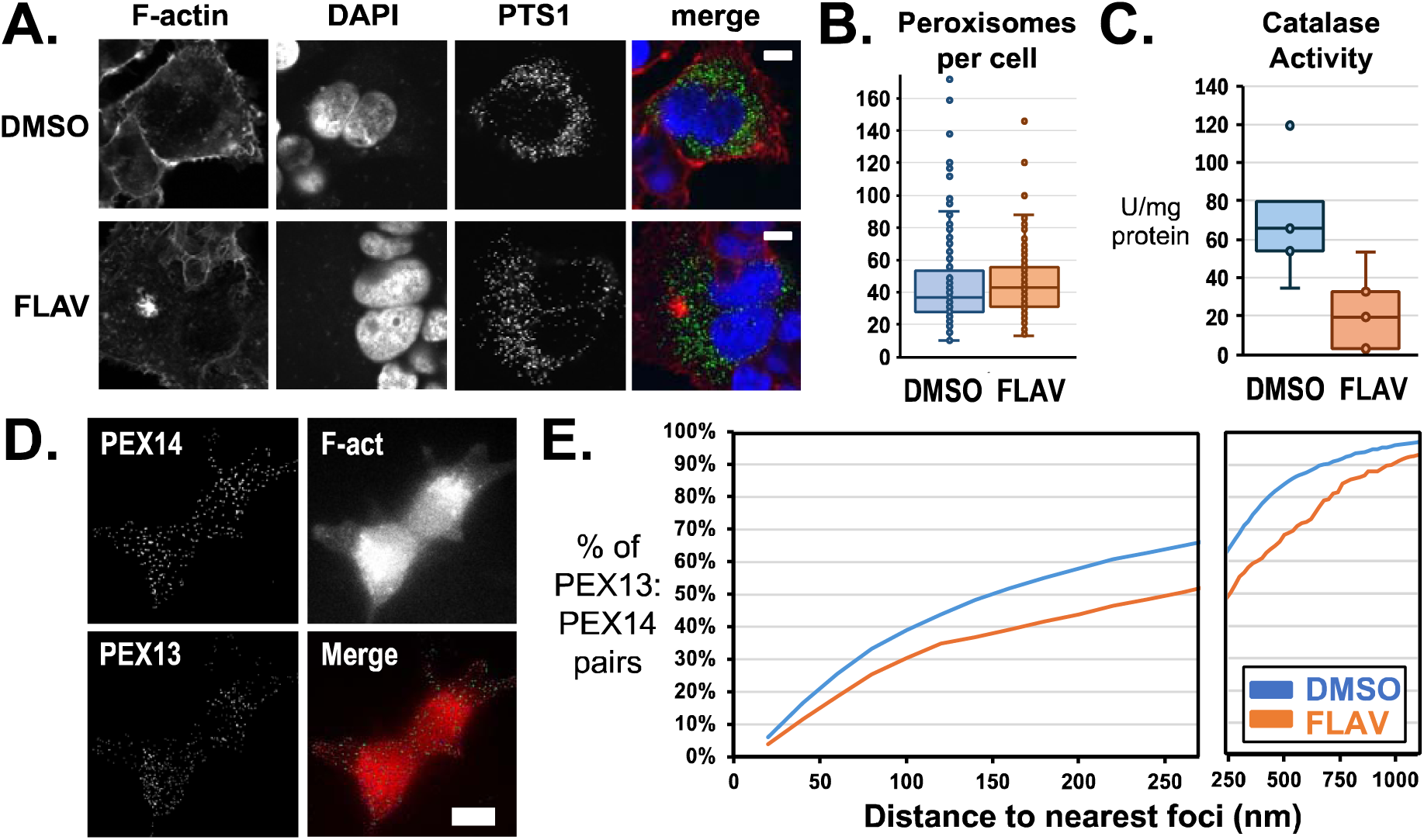
Interactions of PEX13 and PEX14 following treatment of cells by flavopiridol. (**A**) Peroxisomes were detected by imaging of a GFP protein fused to a peroxisome targeting signal, PTS1, for cells treated with 1 µM flavopiridol for 1 hour, FLAV, or vehicle-only, DMSO. (**B**) Number of peroxisomes per cell at 1 hour after flavopiridol treatment was unchanged. (**C**) Catalase activity was significantly reduced by flavopiridol. (**D**) N-STORM microscopy was used to measure the distances between PEX13 and PEX14 proteins. (**E**) The cumulative plot shows the distances measured between PEX13 and the nearest PEX14 neighbor.

As initially noted, membrane bound proteins involved in protein transport featured prominently among peroxisome proteins depleted from particles by flavopiridol treatment. PEX14 was one such protein with a well characterized interaction with PEX13 to form peroxisome pore structures. We confirmed detection of PEX14 and PEX13 by targeted antibodies through confocal immunofluorescent microscopy. Microscopy also revealed flavopiridol induced no change in PEX13 or PEX14 association with peroxisomes (**Supplemental Figure 4**). We used Stochastic Optical Reconstruction Microscopy (STORM) to measure interactions of PEX14 and PEX13 in intact cells (**Figure 5D**). For vehicle treated cells, 50% of PEX14 proteins (N = 2073) were within 150 nm to the nearest PEX13 compared to only 37% for flavopiridol (N = 398) treated cells (p < 10^-14^, **Figure 5E**).

## Discussion

Subcellular structures, such as non-membrane organelles and condensates, play central roles in metabolism, RNA processing, and signaling. In this study, we used crosslinking-assisted SEC-MS as a screening-level approach to survey how large protein assemblies shift across different cell lines and after acute transcriptional inhibition by flavopiridol. This method does not resolve direct protein–protein contacts, but it does reveal crosslink-stabilized association patterns that change with cellular state^24, 26^.

Across the cell lines examined, the two Ewing sarcoma models (A673 and SK-N-MC) showed increased stabilization of splicing factors, microtubule-associated proteins, and cell-division machinery in larger crosslinked structures. In contrast, nucleolar components and membrane-associated proteins from mitochondria and ER appeared less enriched in large fractions relative to the other lines. These observations suggest altered organization of several cellular compartments in these EWS-FLI1 positive backgrounds, though we interpret them cautiously given the long culture histories and diverse origins of the cell lines^36–39^. Our nucleolar measurements, for example, show more nucleoli per nucleus in A673 and in HEK293T cells expressing EWS-FLI1. We do not detect changes in nucleolar area. This pattern is consistent with altered nucleolar organization, though not a defect in nucleolar formation, which has been noted by previous studies of Ewing sarcoma^40, 41^.

Because Ewing sarcoma is driven primarily by the EWS-FLI1 fusion and has few secondary mutations, changes in nuclear architecture are expected. Fusion proteins that drive Ewing Sarcoma, particularly EWS-FLI1, can form condensates in cells that comprise of other nuclear proteins^42–44^. However, the shifts we observe in cytoplasmic and mitochondrial assemblies indicate that the influence of EWS-FLI1 may extend beyond direct nuclear interactions. Future work using isogenic systems or primary/PDX material will be important to determine which of these differences reflect disease-relevant biology versus cell-line-specific features.

Flavopiridol treatment produced rapid and broad changes in the association of transcription and RNA-processing factors with large particles. These results are consistent with our previously published observations that blocking RNA polymerase II disrupts transcriptional condensates, but here we detect a wider set of initiation and processing factors shifting out of large assemblies^24, 25^. Because these changes occur within one hour, they highlight how quickly large macromolecular structures respond to transcriptional stress.

Unexpectedly, flavopiridol also altered peroxisome-associated proteins. We observed reduced catalase activity and increased nanoscale separation between the pore proteins PEX13 and PEX14 by STORM imaging. These findings suggest that transcriptional stress can correlate with peroxisome remodeling, though we interpret this as a drug-specific or stress-linked effect, not a general rule. Follow-up studies using additional transcription inhibitors and washout time courses will be needed to distinguish direct transcriptional consequences from alternative mechanisms, such as off-target or apoptotic responses.

Formaldehyde crosslinking shifts the apparent size distribution of particles, as shown clearly by our no-crosslink controls run under moderately denaturing conditions^23, 24, 26^. For this reason, enrichment values should not be interpreted as stoichiometries. Instead, they reflect crosslink-stabilized co-elution patterns that indicate whether proteins remain associated with large structures under a given condition. Formaldehyde offers a widely used crosslinking chemistry that is well-characterized in many in vivo proteomics studies of protein assemblies^20, 22, 45, 46^. Additional characterization with alternate conditions or crosslinkers such as DSSO, will further strengthen the robustness of these calls in future work^19, 20^.

The SEC fractionation used here has limited resolution and cannot distinguish soluble aggregates from bona fide macromolecular assemblies^19, 22^. We therefore position this approach as a new broad, unbiased screen for structural remodeling that requires orthogonal validation^23, 24^. As illustrated by the peroxisome findings, SEC-MS can highlight unexpected changes, but confirming specific molecular interactions, such as the PEX13–PEX14 separation, still requires targeted assays like microscopy^5, 47^.

Despite these limitations, crosslink-assisted SEC-MS provides a rapid way to detect condition-dependent shifts in large protein assemblies across the cell. Many cellular processes rely on dynamic, multicomponent structures whose organization is still poorly understood. As with the early studies of transcriptional condensates, systematic profiling of assembly-level changes may reveal new regulatory mechanisms and help explain how diseases or therapies reshape local molecular environments. Because protein assemblies can concentrate or exclude drugs and their targets, mapping these structural changes may also provide insight into treatment responses and vulnerabilities^14^.

## Supplemental information

This article contains supplemental data.

## Data availability

The mass spectrometry proteomics data have been deposited to the ProteomeXchange Consortium via the PRIDE partner repository with the dataset identifier PXD067136.

## Acknowledgements

This work was supported by the National Institutes of Health [CA238499 and CA259570 to J.C.S.] and the American Cancer Society [RSG-18-237-01-DMC to J.C.S.]. R.A.V. was also supported by the Initiative to Maximize Student Diversity/NIH program R25 GM062584. Additional support was by the Sloan Indigenous Graduate Partnership, funded by the Sloan Foundation Higher Education Program, to R.A.V. [G-2017-7327] and J.J.A. [G-2020-12601]. J.J.A. was also supported by the National Cancer Institute under award number T32CA009213. M.Z. was supported by 22-1041992-01-DPBACC from the American Cancer Society. This research also was supported by the National Cancer Institute of the National Institutes of Health under award number P30CA023074 through the University of Arizona Comprehensive Cancer Center shared resources.

