## Supplemental Figures for "Global Profiling of Remodeled Subcellular Structures Due to Drug Treatment and Disease"

### Supplemental Figure 1

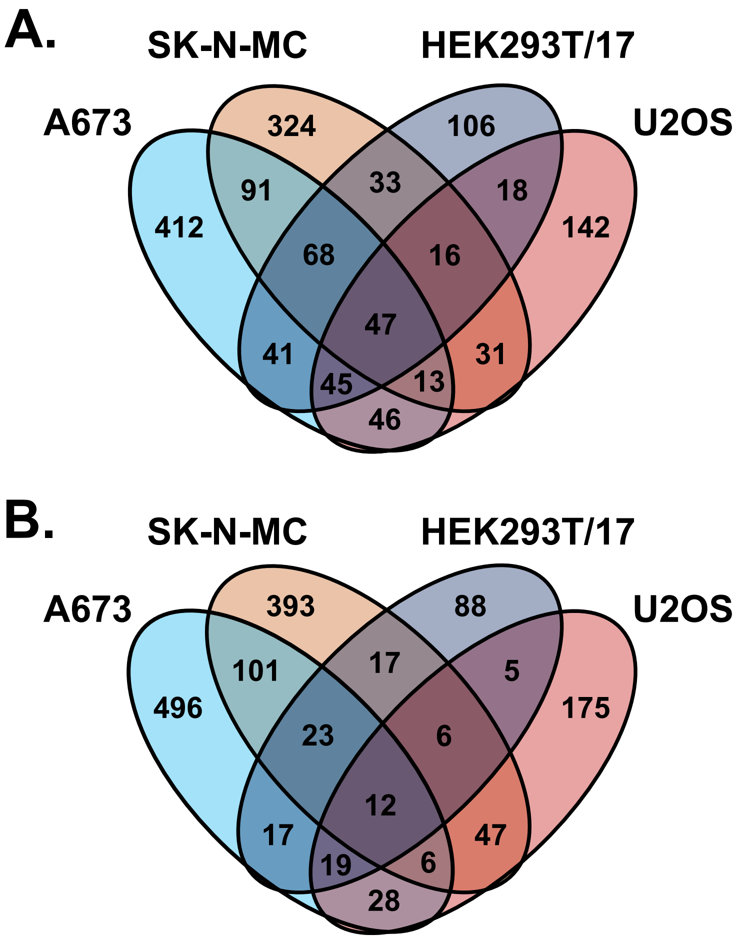


**Supplemental Figure 1:** Number of enriched proteins shared or unique to Ewing or non-Ewing sarcoma cell lines for (**A**) large-sized or (**B**) medium-sized particle pools.

### Supplemental Figure 2

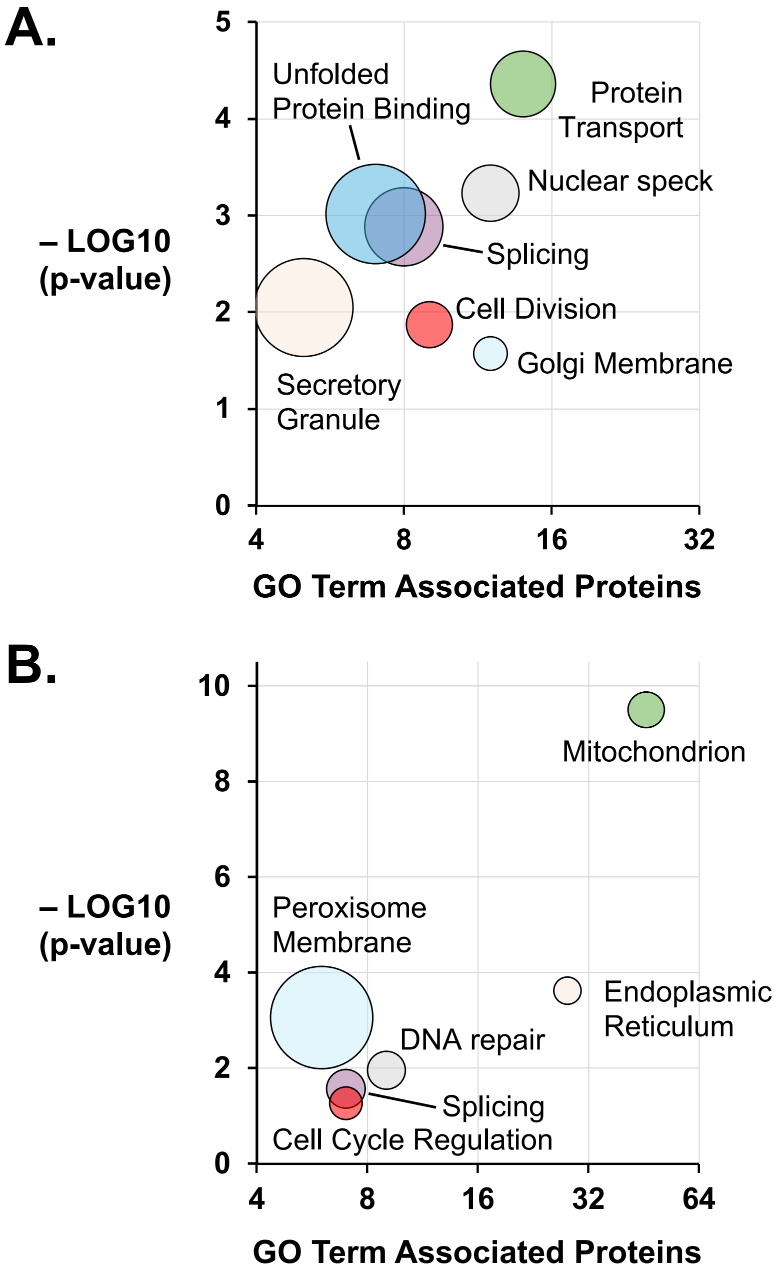


**Supplemental Figure 2:** Bubble plots show functional annotation clusters and GO terms for proteins with (A) increased enrichment by crosslinking in large or medium particles for cells treated by flavopiridol or (B) decreased in enrichment. Diameters indicate fold enrichment of associated proteins.

### Supplemental Figure 3

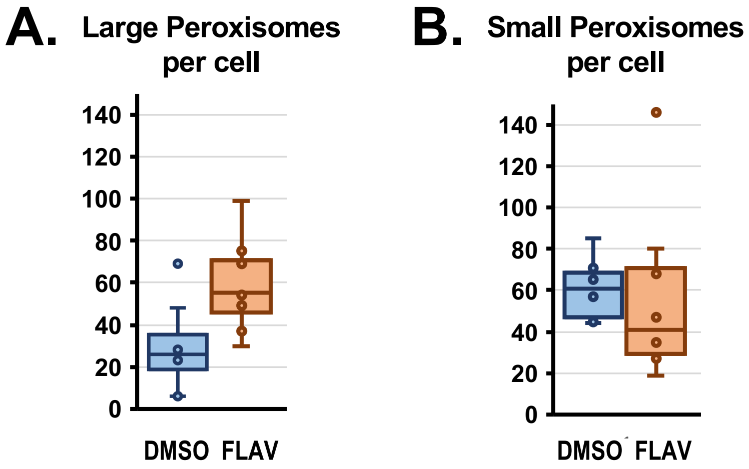


**Supplemental Figure 3:** Numbers of (**A**) large or (**B**) small peroxisomes in cells treated by either vehicle (DMSO) or flavopiridol. The difference in large particles is significant ($p = 0.017$).

### Supplemental Figure 4

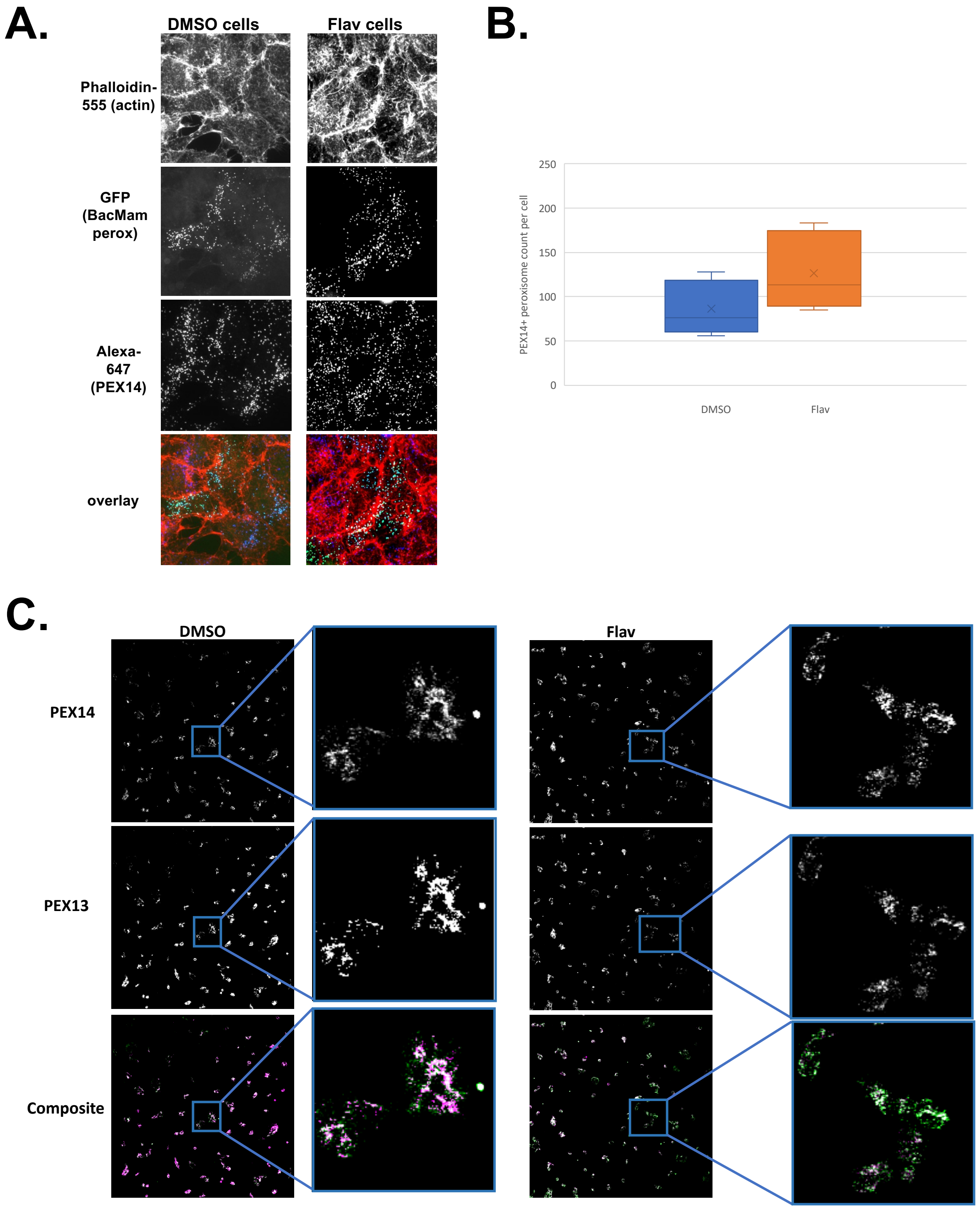


**Supplemental Figure 4:** Number of peroxisomes (BacMam perox) that colocalize with PEX14 (**A**) in Sora confocal microscopy is not significantly different (**B**) between DMSO or flavopiridol treated cells. (**C**) The number of PEX14 and PEX13 detected by STORM microscopy is all not different.
